## Supplementary material for "Clp protease and antisense RNA jointly regulate the global regulator CarD to mediate mycobacterial starvation response": This file includes seven supplementary figures and three supplementary tables.

This PDF file includes:

1. Seven supplementary figures
2. Three supplementary tables

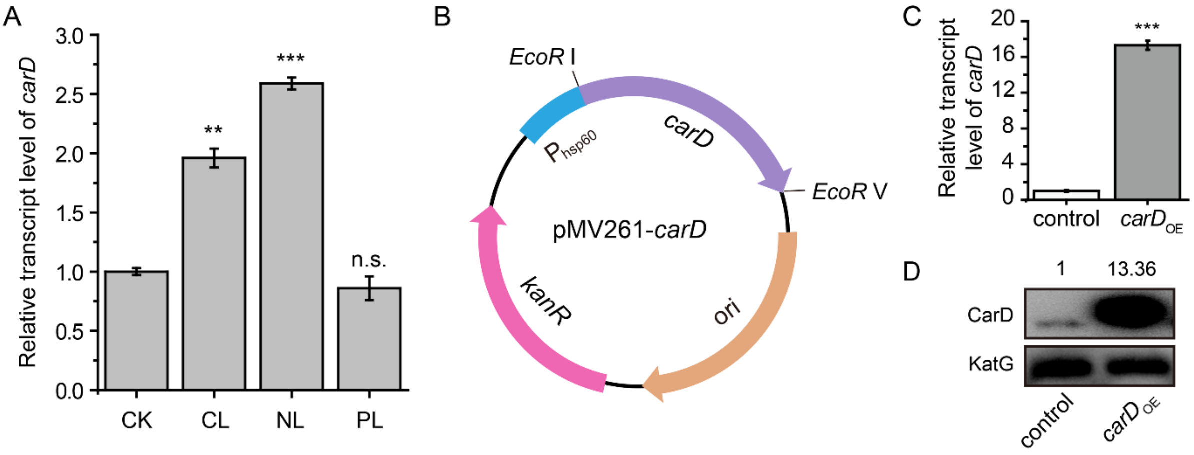


**Figure 1–figure supplement 1. Changes of *carD* levels in *M. smegmatis* under different conditions and different strains. A**, *carD* mRNA levels under different starvation conditions. CK represents untreated exponential cells; CL, NL, and PL represent exponential cells transferred into carbon-limiting, nitrogen-limiting, and phosphorus-limiting media for 4 h, respectively. Statistical analysis was done using Student's t-test, with ** indicating p-value <0.005, *** indicating p-value <0.001, and n.s. indicating p-value >0.05. Error bars indicate the standard deviation of three biological replicates. **B**, Schematic diagram for the construction of the *carD* overexpression plasmid. The coding sequence of *carD* was cloned into multiple-copy plasmid pMV261 between the *Eco*R I and *Eco*R V restriction sites, which allowed *carD* to be transcribed from the *hsp60* promoter on the plasmid. **C** and **D**, the mRNA and protein levels of *carD* in the control and *carD* overexpression strains (*carD*$_{\mathrm{OE}}$), respectively. The number on each band of the Western blot results represent their relative quantitative values, which are normalized with respect to their corresponding loading controls.

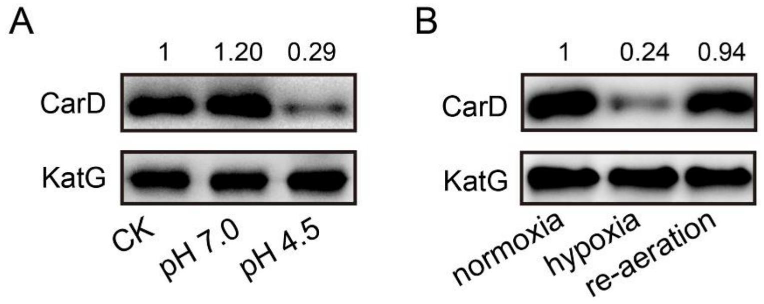


**Figure 2–figure supplement 1. Changes of CarD levels in *M. smegmatis* under host-like stress conditions.** **A**, CarD protein levels in mc$^2$155 under different pH conditions. CK indicates the untreated exponential cells; pH 7.0 and pH 4.5 indicate the exponential cells transferred into the media with corresponding pH values for 4 h. **B**, CarD protein levels in mc$^2$155 under different oxygen availability conditions. The number on each band of the Western blot results represent their relative quantitative values, which are normalized with respect to their corresponding loading controls.

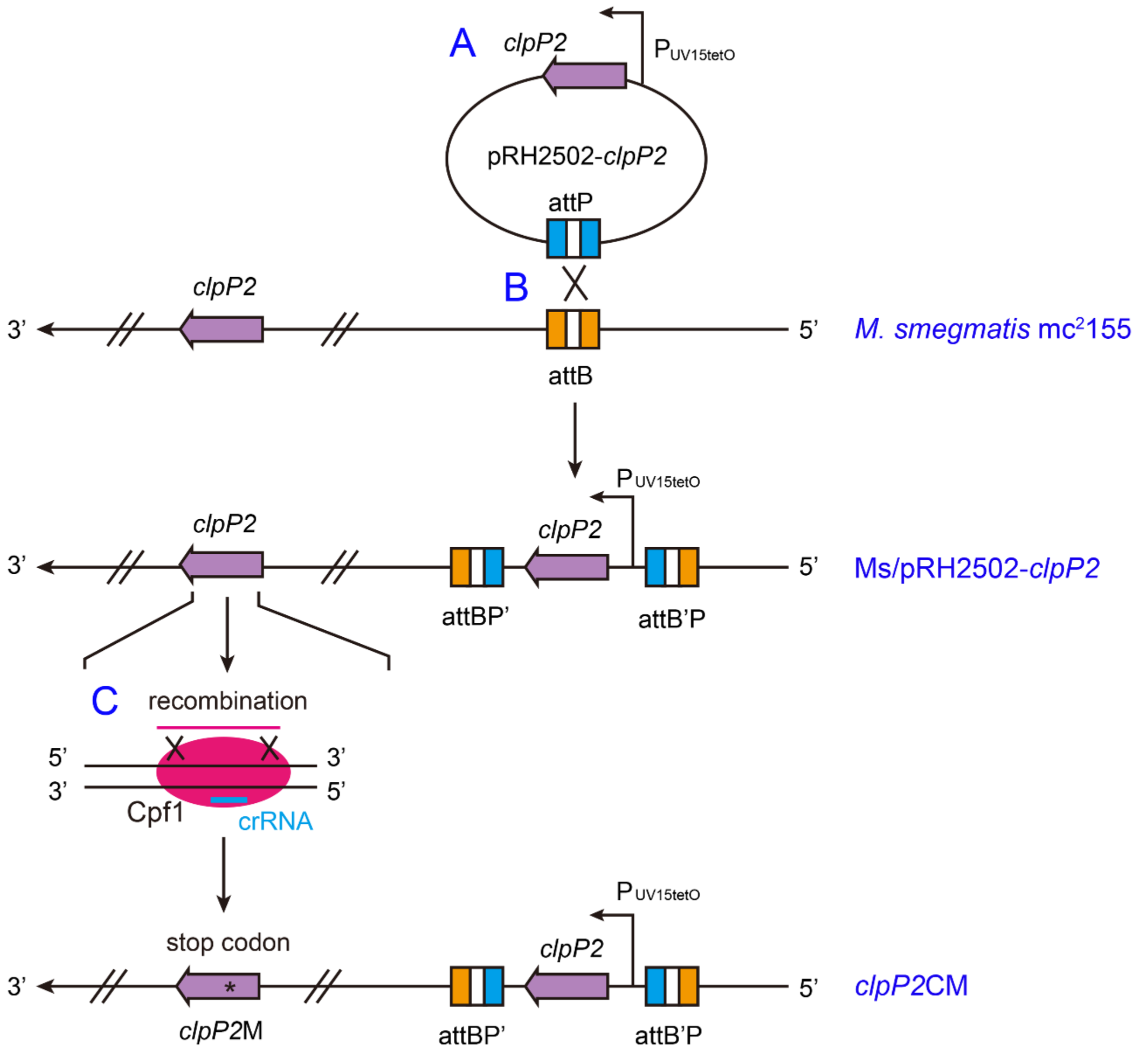


**Figure 3–figure supplement 1. Schematic diagram for the construction of the *clpP2* conditional mutant. A,** *clpP2* gene amplified with *clpP2*-F/R primer pair (Supplementary File 3) was ligated to the pRH2502 integration plasmid to obtain pRH2502-*clpP2* recombinant plasmid, in which *clpP2* is under the control of ATc-inducible promoter $P_{UV15tetO}$. **B,** the pRH2502-*clpP2* plasmid was transformed and integrated into $mc^2155$ genome by *attB*-*attP* mediated site-specific recombination, to obtain Ms/pRH2502-*clpP2* strain. **C,** CRISPR/Cpf1-mediated mutagenesis was used for the mutation of the endogenous *clpP2* gene to obtain the *clpP2* conditional mutant *clpP2CM*.

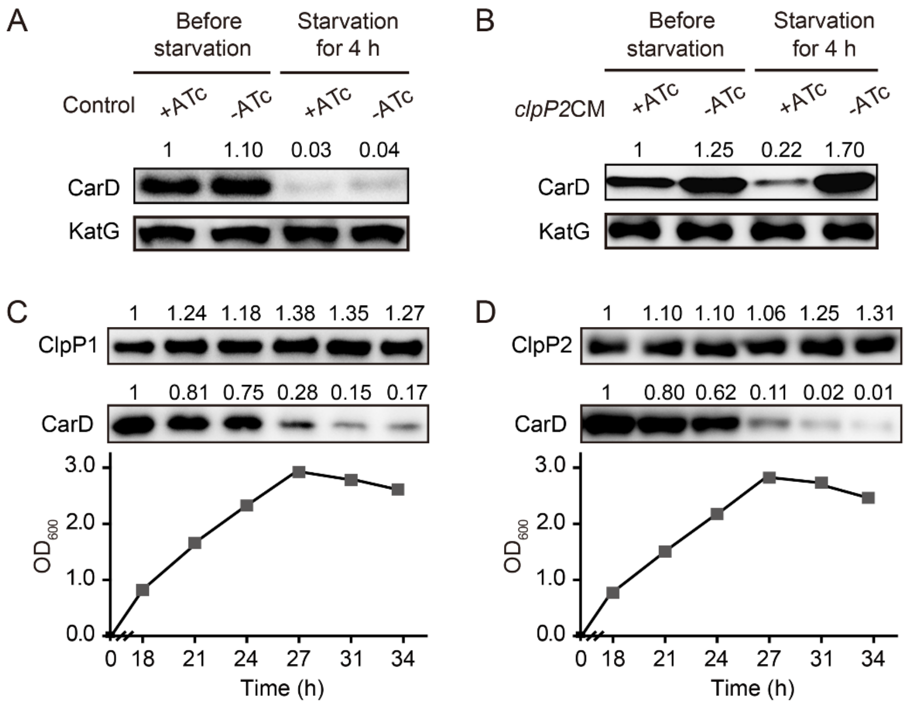


**Figure 3–figure supplement 2. Clp protease degrades CarD under the starvation condition. A** and **B,** the starvation experiments on control and *clpP2CM* cells, respectively. The cells used for starvation were harvested at the exponential phase (three hours before the stationary phase). **C** and **D**, protein levels of ClpP1 and ClpP2, respectively, at different time points in *M. smegmatis* cells. For all panels, the number on each band of the Western blot results represent their relative quantitative values.

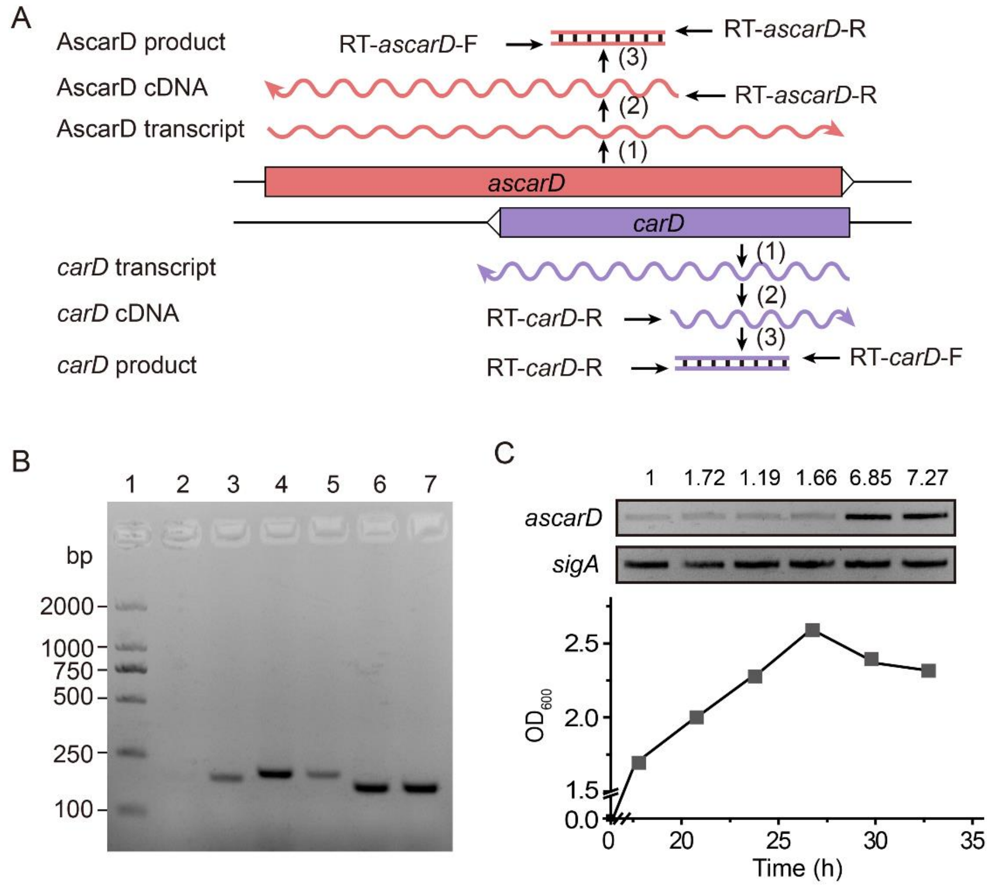


**Figure 4–figure supplement 1. RT-PCR analysis of the transcriptional levels of *ascarD* and *carD*. A,** schematic diagram of the strand-specific RT-PCR. Step (1) represents transcription of the *ascarD* and *carD* genes; step (2) represents *ascarD* and *carD* transcripts reverse transcribed into the corresponding cDNAs with RT-*ascarD*-R/RT-*carD*-R primers (Supplementary File 3); step (3) represents amplification of AscarD and *carD* cDNA with RT-*ascarD*-F/R or RT-*carD*-F/R primer pairs (Supplementary File 3), respectively. **B,** the RT-PCR results at different growth phases. Lane 1 is the DL2000 ladder marker, lanes 2 and 3 show the *ascarD* RNA levels at MEP and MSP, respectively; lanes 4 and 5 show *carD* RNA levels at MEP and MSP, respectively; lanes 6 and 7 show the RNA levels of internal reference gene *sigA* at MEP and MSP, respectively. **C,** *ascarD* RNA levels throughout the growth phase as measured by RT-PCR; *sigA* was used as an internal reference gene. The number on each band of the RT-PCR results represent their relative quantitative values, which are normalized with respect to their corresponding loading controls.

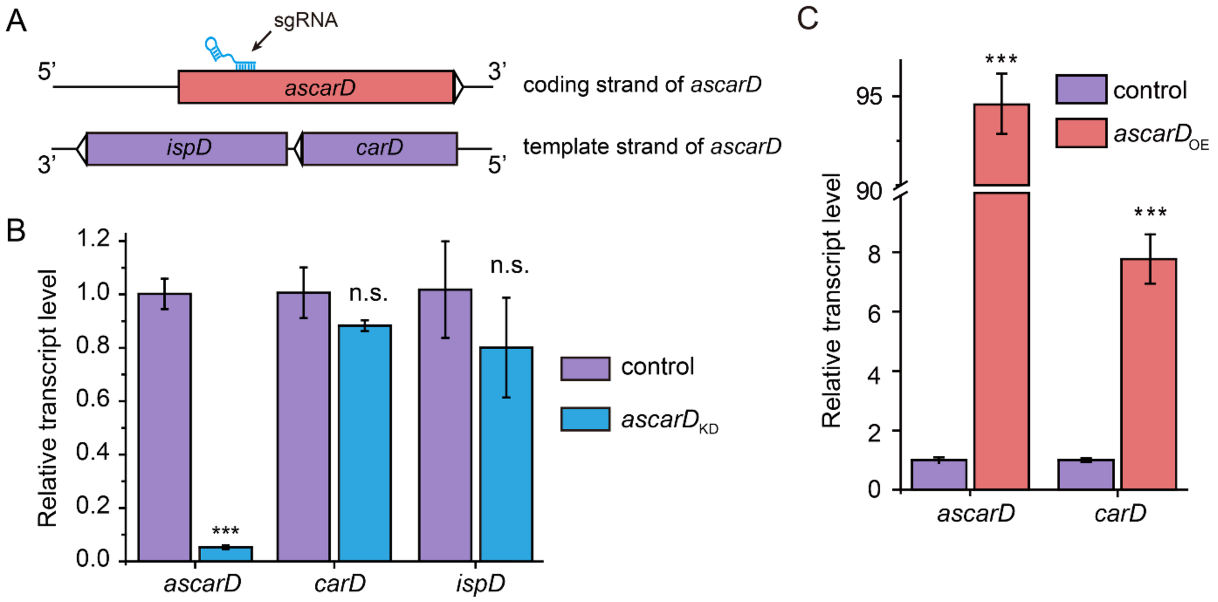


**Figure 5–figure supplement 1. The expression levels of *carD* and *ascarD* in different strains. A,** Schematic diagram for the location of sgRNA targeting *ascarD*. **B**, Transcript levels of *ascarD*, *carD*, and *ispD* in *ascarD*$_{\mathrm{KD}}$ and control strains. The mc$^2$155 strain transformed with pRH2521 empty vector was used as the control. **C**, transcript levels of *ascarD* and *carD* in *ascarD*$_{\mathrm{OE}}$ and control strains. The mc$^2$155 strain transformed with pMV261 empty vector was used as the control. *sigA* was used as the internal reference gene of qRT-PCR. Error bars indicate the standard deviation of three biological replicates. Statistical testing was done using the Student's t-test, with *** indicating p-value <0.001, n.s. indicating p-value >0.05.

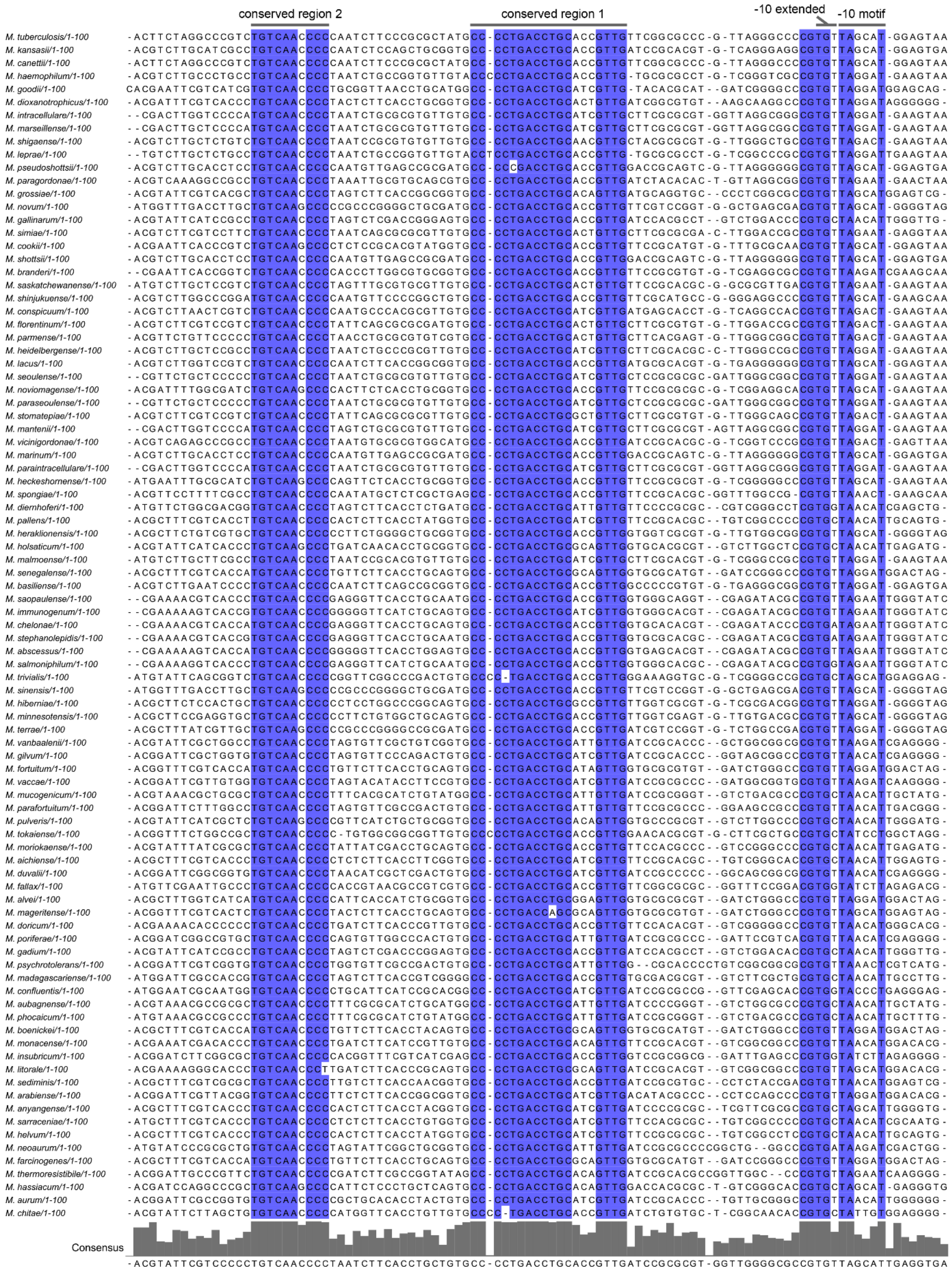


**Figure 7–figure supplement 1. Alignment of mycobacterial *carD* promoter sequences.** The 100 bp promoter sequences of *carD* from 91 different mycobacteria were aligned by Clustal W, and the highly conserved nucleotides were marked in blue.

**Supplementary File 1 The top 20 up-regulated genes in the CarD$^{K125A}$ mutant**

| Gene[a] | Fold change[b] | Protein family[c] | Function[c] |
|---|---|---|---|
| *Rv2623* | 36.03 | pfam00582 | Universal stress protein |
| *Rv2624c* | 25.19 | pfam00582 | Universal stress protein |
| *Rv3129* | 19.46 | pfam12900 | Pyridoxamine 5'-phosphate oxidase family protein |
| *Rv2626c* | 17.49 | pfam00571 | Hypoxic response protein 1, CBS domain pair |
| *Rv3128c* | 15.14 | N/A | DDE-type integrase/transposase/recombinase |
| *Rv2030c* | 14.41 | pfam05139 | Erythromycin esterase family protein |
| *Rv2007c* | 11.51 | COG1146 | Ferredoxin FdxA |
| *Rv2029c* | 11.42 | COG1105 | 6-Phosphofructokinase PfkB |
| *Rv2625c* | 11.39 | COG1994 | Putative zinc metalloprotease Rip3 |
| *Rv2627c* | 10.95 | N/A | Alpha/beta hydrolase |
| *Rv0022c* | 9.31 | pfam02467 | Transcriptional regulator WhiB5 |
| *Rv2031c* | 7.21 | pfam00011 | Heat shock protein HspX |
| *Rv1736c* | 6.74 | COG5013 | Nitrate reductase NarX |
| *Rv1738* | 5.89 | pfam08962 | Bacterial hibernation-promoting factor (PDB: 4wpy) |
| *Rv2628* | 5.66 | N/A | Hypothetical protein |
| *Rv1997* | 5.56 | COG0474 | Cation-transporting ATPase CtpF |
| *Rv2028c* | 5.52 | pfam00582 | Universal stress protein |
| *Rv1130* | 5.06 | COG2079 | 2-methylcitrate dehydratase PrpD |
| *Rv1733c* | 4.87 | N/A | Transmembrane protein |
| *Rv1737c* | 4.62 | COG2223 | Nitrate/nitrite transporter NarK2 |

[a]- The underlined genes belong to the dormancy regulon identified previously.

[b]- Change in gene expression in the CarD$^{K125A}$ mutant compared to the wild-type strain, based on the previously published RNA-seq data (GEO accession number: GSE131043).

[c] - Protein family assignments in Pfam and COG databases, where available. Functional assignments are from GenBank and/or RefSeq.

**Supplementary File 2 Strains used in this study**

| Strain | Characteristics used in this work | Source |
|---|---|---|
| $mc^2155$ | Wild-type strain of *M. smegmatis* $mc^2155$, abbreviated as Ms | Lab stock |
| BCG | Wild-type strain of *M. bovis* BCG | Lab stock |
| H37Ra | Wild-type strain of *M. tuberculosis* H37Ra | Lab stock |
| Ms/pMV261 | $mc^2155$ with pMV261 empty vector | This study |
| $ascarD_{OE}$ | *ascarD* overexpression strain | This study |
| Ms/pRH2521 | $mc^2155$ with pRH2521 empty vector | This study |
| $ascarD_{KD}$ | *ascarD* knock-down strain | This study |
| $carD_{OE}$ | *carD* overexpression strain | This study |
| Ms/pRH2502-*clpP2* | $mc^2155$ strain integrated with pRH2502-*clpP2* plasmid | This study |
| *clpP2*CM | *clpP2* conditional mutant | This study |
| AAAS_del | CarD C-terminal “AAAS” motif deletion mutant | This study |
| Ms/pRH2502-*clpC1*-His | used for the ClpC1 protein level determination | This study |
| Δ*sigF* | $mc^2155$ with *sigF* knocked out | This study |
| Ms/pMV261-P*ascarD*-*lacZ* | used for the promoter activity determination of *ascarD* in $mc^2155$ strain | This study |
| Δ*sigF*/pMV261-P*ascarD*-*lacZ* | used for the promoter activity determination of *ascarD* in Δ*sigF* strain | This study |
| Ms/$P_{PUCP}$ | used for the promoter activity determination of $P_{PUCP}$ in $mc^2155$ strain | This study |
| Ms/$P_{PUCP*}$ | used for the promoter activity determination of $P_{PUCP*}$ in $mc^2155$ strain | This study |
| AsM | *ascarD* promoter mutant, with the -10 motif changed from “GGGTA” to “**C**GG**C**A” | This study |
| AsM/AAAS_del | Double mutant of the *ascarD* promoter and CarD “AAAS” motif | This study |

### Supplementary File 3 Oligonucleotides used in this study

| Primer name | Use in this work | Sequence 5′–3′ |
|---|---|---|
| RT-*ascarD*-F | Forward primer for AscarD detection by RT-PCR | TCTCGGCCTTGGCGTCATC |
| RT-*ascarD*-R | Reverse primer for AscarD detection by RT-PCR | GTCGCGTCGCTACAAGGC |
| RT-*carD*-F | Forward primer for *carD* transcript detection by RT-PCR | TCGTCTATCCACACCACGGTGC |
| RT-*carD*-R | Reverse primer for *carD* transcript detection by RT-PCR | TCGGCTCCTCGGTGTGCG |
| RT-*ispD*-F | Forward primer for *ispD* transcript detection by RT-PCR | CAGACGCCTCAGGGTTTCCA |
| RT-*ispD*-R | Reverse primer for *ispD* transcript detection by RT-PCR | ATCTTGAATGCCAGCGGGTC |
| RT-*sigA*-F | Forward primer for *sigA* detection by RT-PCR | CGTTCCTCGACCTCATCCAG |
| RT-*sigA*-R | Reverse primer for *sigA* transcript detection by RT-PCR | TGATCACCTCGACCATGTGC |
| *carD*$_{OE}$-F | Forward primer for *carD* overexpression | CCGGAATTCATTTTTAAGGTCGGAGACAC CGTC |
| *carD*$_{OE}$-R | Reverse primer for *carD* overexpression | ATAGATATCGGACGCGGCGGCCAAAACC TCGT |
| *clpP2*-F | Forward primer for *clpP2* amplification | gcgaaggagatatacatatgATGAGCAACATaCATCC GTC |
| *clpP2*-R | Reverse primer for *clpP2* amplification | gttccgagtttgttaattaaACCCGTTGTCGTGTCAGA |
| *clpP2*-crRNA-F | Forward primer of *clpP2*-crRNA | ATATCCGTCACTGGACGCCCGGCTGCAA |
| *clpP2*-crRNA-R | Reverse primer of *clpP2*-crRNA | AGCTTTGCAGCCGGGCGTCCAGTGACGGA TATCT |
| *clpP2*-lag | 59 nt lag sequence (targeting the lagging strand of DNA replication), used for *clpP2* mutation | GACGGCAGGATGTAGCGGGCCTGCGGCTtt aCGGATGAATGTTGCTCATTTGTCTAGTC |
| AAAS-crRNA-F | Forward primer of AAAS-crRNA | ATTGGCCGCCGCGTCCTGATCTGTTCGA |
| AAAS-crRNA-R | Reverse primer of AAAS-crRNA | AGCTTCGAACAGATCAGGACGCGGCGGC CAATCT |

| | | |
|---|---|---|
| AAAS-lag | 79 nt lag sequence, used for CarD AAAS motif deletion | ACGGCAACCGTCGCCACCGTTTACGTCCCGAACAGATCACAAAACCTCGTCGAGGATGGTCTCGGCCTTGGCGTCATCG |
| *clpC1*-F | Forward primer for *clpC1*-His construction | atctagacatcatcggtaccCGCAACGGACACAATGTGTCGA |
| *clpC1*-R | Reverse primer for *clpC1*-His construction | gttccgagtttgttaattagtggtggtggtggtggtgCTCCGTGCCTGCGGTCGG |
| 5'-RACE-*ascarD*-F | Forward primer for AscarD TSS identification | GACCACGCGTATCGATGTCGACTTTTTTTTTTTTTTTTV |
| 5'-RACE-*ascarD*-R | Reverse primer for AscarD TSS identification | CCCGCGCTCACCGACGAGTCGAAAC |
| *sigF*-crRNA-F | Forward primer of *sigF*-crRNA | ATCGCGACAACAGCTGGTCGGTGAAGGA |
| *sigF*-crRNA-R | Reverse primer of *sigF*-crRNA | AGCTTCCTTCACCGACCAGCTGTTGTCGCGATCT |
| *sigF*-lag | 59 nt lag sequence, used for *sigF* mutation | GGAGTTCCTTGAGCCGGCGAGGCACCTTCAGTGCCTGCGGACCTCGCCCATGATCGTGG |
| *ascarD*-sgRNA-F | Forward primer of *ascarD*-sgRNA | GGGAGCGCTCGCTCTCGAAGCGGC |
| *ascarD*-sgRNA-R | Reverse primer of *ascarD*-sgRNA | AAACGCCGCTTCGAGAGCGAGCGC |
| *ascarD*$_{OE}$-F | Forward primer for *ascarD* overexpression | GCTCTAGACCAGGCACAGCAGACGACAC |
| *ascarD*$_{OE}$-R | Reverse primer for *ascarD* overexpression | CCCAAGCTTTGATCTGGGCCCGTGTTAGG |
| P*ascarD-lacZ* -F | Forward primer for the construction of *ascarD* promoter reporting plasmid | CGGTACCAGATCTTTAAATCTAGACCAGGCACAGCAGACGACACGG |
| P*ascarD-lacZ* -R | Reverse primer for the construction of *ascarD* promoter reporting plasmid | AAACGACGGGATCCATTGAAGCTTCCGGGTGGTGGCTGCGCTGA |
| PUCP-F | Forward primer for PUCP construction | CGGTACCAGATCTTTAAATCTAGATGGCGATCATCAGGGGAATGT |
| PUCP-R | Reverse primer for PUCP construction | AAACGACGGGATCCATTGAAGCTTGGATGGTCTCGGCCTTGGCG |
| AsM-crRNA-F | Forward primer of *ascarD*-crRNA | ATCGAGCACCGCGCCGTTGGCGTCGACA |
| AsM-crRNA-R | Reverse primer of *ascarD*-crRNA | AGCTTGTCGACGCCAACGGCGCGGTGCTCGATCT |
| AsM-upstream-F | Forward primer of the *ascarD* upstream | GTCGAATTCCAGGCACAGCA |

| | | |
|---|---|---|
| AsM-upstream-R | Reverse primer of the *ascarD* upstream | TACGCCCGAGCGGGCCGGTCTGCGCGCTGTCCAG |
| AsM-downstream-F | Forward primer of the *ascarD* downstream | CCGGCCCGCTCGGGCGTACCGAGCACCGCGCCGTTGGCGTCGACAGCCTTT |
| AsM-downstream-R | Reverse primer of the *ascarD* downstream | GAAAGCATTCGTGACACTGGGG |
